# A SENSORIZED 3D-PRINTED KNEE TEST RIG FOR EXPERIMENTAL VALIDATION OF PATELLAR TRACKING AND CONTACT SIMULATION AFTER TOTAL KNEE REPLACEMENT

**DOI:** 10.1101/2024.02.13.580095

**Authors:** Florian Michaud, Francisco Mouzo, Daniel Dopico, Javier Cuadrado

## Abstract

Total knee arthroplasty aims to relieve pain and restore function in the affected joint through artificial implants. Despite advances in design and surgical techniques, there are complications associated with surgery, especially concerning the patella. The use of musculoskeletal models in orthopedic surgery allows for objective prediction of postoperative function and optimization of results for each patient. To ensure that simulations are trustworthy and can be used for predictive purposes, comparing simulation results with experimental data is crucial. Although progress has been made in obtaining 3D bone geometry and estimating contact forces, validation of these predictions has been limited due to the lack of direct in vivo measurements and the economic and ethical constraints associated with available alternatives. In this study, an existing commercial surgical training station was transformed into a sensorized test bench. The use of 3D-printed models and sensors allowed low-cost and reproducible experimental validation of computer simulation (patellar movement and forces) while avoiding ethical issues.

## 1. Introduction

Aims of Total Knee Replacement (TKR) are to alleviate pain, restore function, and lead to an improved quality of life (Tekin, Ünver, and Karatosun 2012). Despite the continuous advances in implant design and surgical techniques, numerous TKR complications are still observed, the 10% of them being associated to patellar complications which may require repetition of surgical procedures (Putman et al. 2019). To avoid extensor mechanism complications and ensure good functional outcome, obtaining proper patellar tracking is one of the most important goals of TKR. Poor patellar tracking can result in increased postoperative contact pressures, patellar tilt, patella subluxation, or dislocation (Goyal, Matar, and Parvizi 2012). Use of musculoskeletal models for orthopedic surgeries offers an objective prediction of post-treatment function and allows clinicians to explore different treatment options, to reduce the level of subjectivity in the treatment planning process, and to optimize clinical outcomes on an individual patient basis (Fregly et al. 2012). However, while the potential of the method is great, the research community has made only limited progress at validating its predictions due to the lack of direct in vivo measurements (Fregly 2021), which are limited for technological and ethical reasons. By comparing simulation results with experimental data, researchers can gauge the model’s fidelity and reliability. This is crucial for ensuring that simulations are trustworthy and can be used for predictive purposes. It is essential for surgeons, as well as for the scientific community, to have confidence that the computer simulation accurately represents real-world phenomena.

Historically, to avoid invasive in vivo measurements in humans, cadaveric data is used to provide generic model information, leading to challenges related to scalability, limits of practical applicability, cost, and ethical compliance (Fischer et al. 2019; Tesfaye et al. 2021). Over the last few decades, thanks to technological advances, some research groups have come to perform some internal in vivo measurements, such as 3D bone geometry from medical images (Stephen et al. 2021), joint kinematics through fluoroscopy (Stiehl et al. 1995), contact forces between femur and tibia by means of instrumented implants (Stansfield et al. 2003; Fregly et al. 2012; D’Lima et al. 2005; Kutzner et al. 2010; Taylor et al. 2017) or, more recently, tendon forces with shear wave tensiometers (Martin et al. 2018). Although some initiatives in the research community allow the sharing of certain in vivo measurements (Fregly et al. 2012), the reality is that few research groups possess the financial, experimental and clinical resources necessary to generate significant new advances in the field.

On the other hand, knee simulators have revolutionized the way researchers, surgeons, and manufacturers approach implant design, durability assessment, surgical techniques, and patient safety, by providing a controlled and repeatable environment for evaluating TKR implants. While the first knee simulator described by Dowson in 1977 was a mechanical device, later, cadaveric knee simulators have emerged as a response to the need for gaining deeper insights into the workings of both the native knee and TKRs (Maag et al. 2021). The underlying concept involves the integration of implants into these cadaveric simulators, with the goal of prescribing specific muscle forces and displacements within the simulator. These controlled inputs serve to replicate both the motion and loads experienced by the implant. Researchers have devised various approaches to achieve this objective, with universities like Oxford, Kansas, and Purdue developing comparable systems for this purpose (Maletsky and Hillberry 2005; Halloran et al. 2010). However, once again, the reality is that few research groups possess the financial, experimental and clinical resources necessary to reproduce these experiments.

In this work, authors aim to achieve low-cost and reproducible experimental validation of computer simulations for patellar trajectory after TKR while avoiding ethical issues by utilizing 3D-printed models and sensors. For this purpose, a commercial training station for knee ligament release which recreates a human leg (Mita n.d.) was adapted. This tool consists of articulated metal supports for the hip and foot, and replaceable inserts for the knee joint. In order to obtain a virtual replica of the system, known as a digital twin, the geometries of the implants were virtually applied and the resulting cut bones were 3D-printed, with real tibia and femur implants being placed in the respective physical bone models. A prosthetic patellar button was attached to a pressure sensor, connected to the tibia via a spring, and to the femur via another spring with lower stiffness. The springs represent the patellar tendon and the quadriceps tendon, respectively, and were connected in series to respective load cells to measure their tensions. The movements of femur, tibia and patella were recorded by an optical motion capture system. The recorded movements of femur and tibia were used as inputs for the simulation. The recorded movement of the patella was used, on the one hand, to approximate its initial position in the simulation, and, on the other hand, in combination with the forces recorded by the pressure sensor and load cells, to validate the results obtained from the simulation, which was performed using a custom-developed library (Dopico 2016; Dopico et al. 2019).

## 2. Material and methods

### 2.1. Test Bench

The commercial training station (Mita Collateral Ligament Release Workstation, Bristol, United Kingdom) for knee ligament release (Mita n.d.), shown in Figure 1, was adapted for this work. The existing commercial structure that recreates a human leg (base with spherical joint for the hip, black metal supports for bones, and a polypropylene foot) was utilized. The replaceable knee inserts were replaced with 3D-printed bones incorporating their respective implants, and several sensors were added.

**Figure 1:**
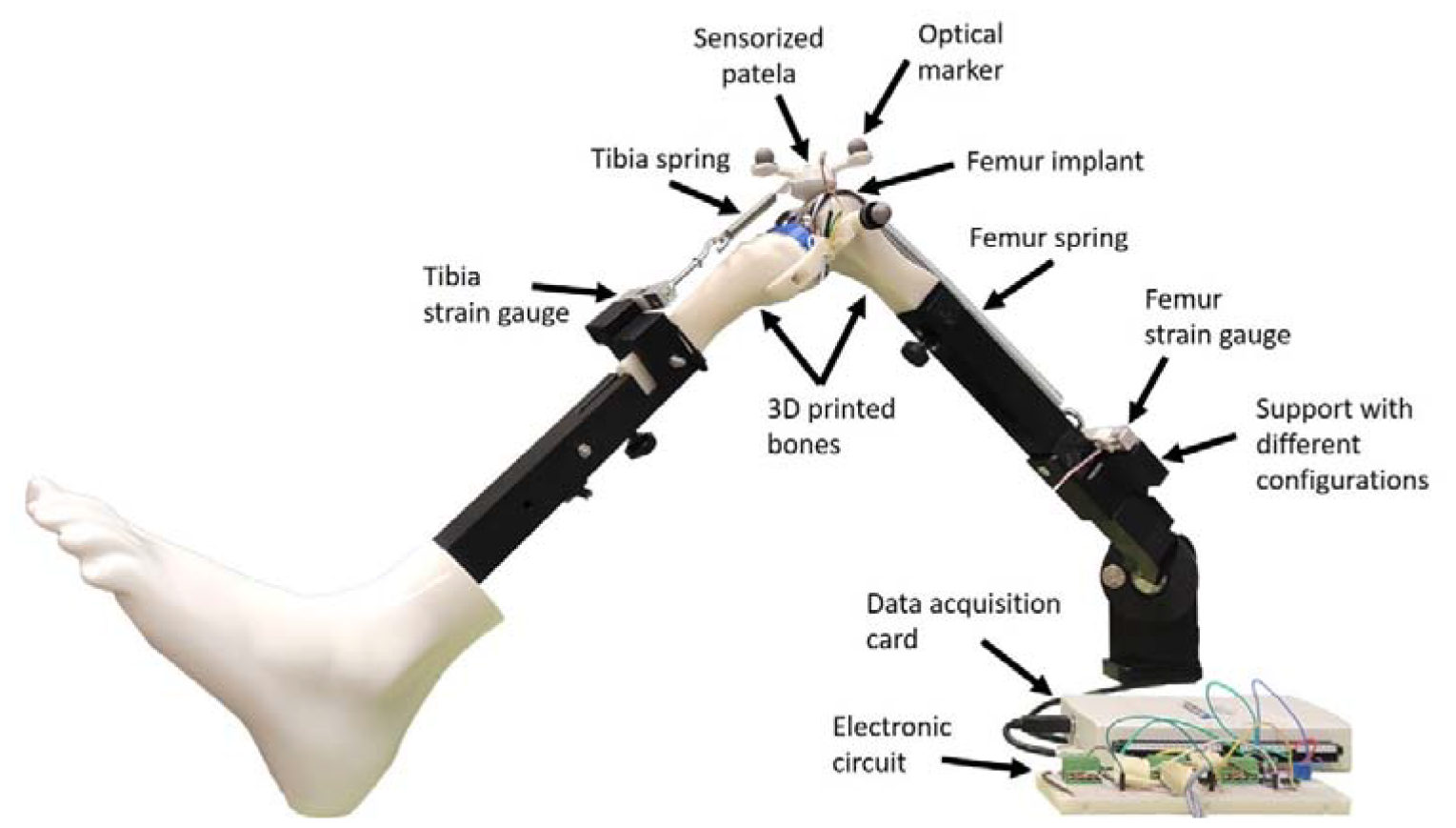
Sensorized test bench.

From the medical images of a patient, it is possible to obtain customized CAD models of their bones. Treatments can be applied virtually to these models, thus simulating their effects. On the other hand, 3D printing allows for the production of physical models of these bones with the applied treatment, resulting in two twin models: the digital model and the physical model. In this work, the case of TKR was addressed. Therefore, the virtual geometries of the implants were applied, and the resulting cut bones were 3D-printed (Prusa I3 MK3S, Prague, Czech Republic), with commercial implants (Microport®) of tibia and femur being placed in their respective bone models.

Motion and force sensors enable the reproduction of movement in the virtual model, adjustment of simulation parameters, and experimental validation of the results. The movements of the femur, tibia, and patella were recorded by an optical motion capture system. Six optical markers were placed to capture the movement of the three bodies constituting the system (two additional markers were used to determine the hip center, which was fixed, through a calibration capture). Due to the preliminary nature of this work, the cruciate ligaments were released (as it happens in the surgery), and the collateral ligaments were treated as rigid bodies (to avoid contact between femur and tibia implants), allowing only one degree of freedom at the knee, in addition to the three rotations at the hip. The patella, on the other hand, was considered a fully free body, in contact with the femoral implant, attached to the tibia via one spring (Figure 1, tibia spring), and to the femur via another spring with lower stiffness (Figure 1, femur spring). To recreate different patellar tracking, the 3D-printed support of the femur strain gauge allowed two different configurations.

The springs represent the patellar tendon and the quadriceps tendon, respectively, and were connected in series with load cells to measure their tensions (Figure 1). For the patella, as seen in Figure 2, a prosthetic patellar button was attached to a pressure sensor through 3D-printed supports onto which three optical markers were fixed. In order to mimic the lubricating effect of synovial fluid in the joint, lubricant was applied to the contacting surfaces.

**Figure 2:**
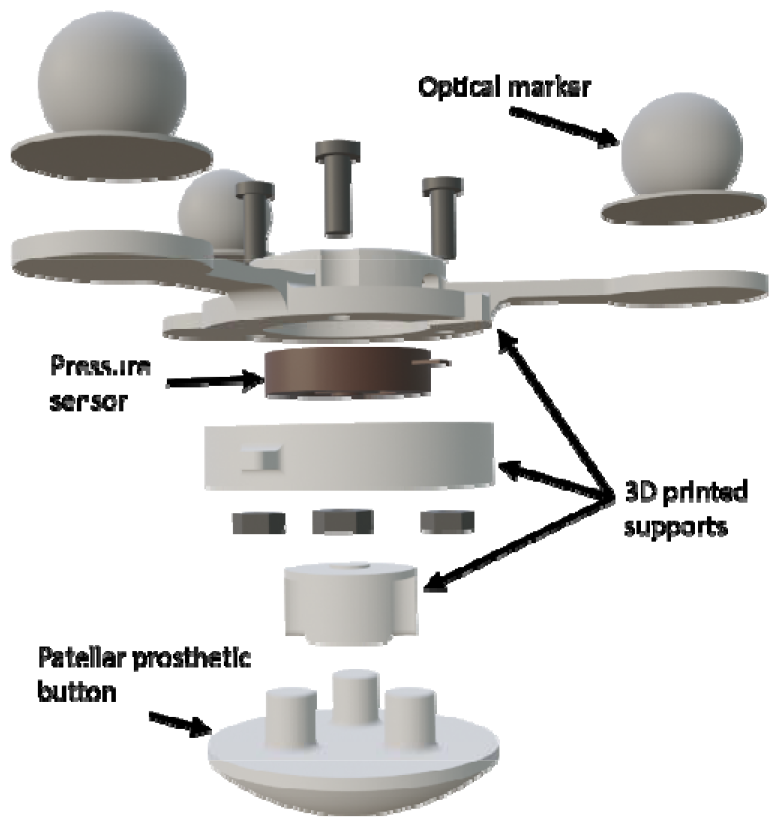
Sensorized patella.

### 2.2. Movement and Experimental Data Collection

The patellar trajectory is defined as the movement of the kneecap relative to the femoral groove during knee flexion and extension (Katchburian et al. 2003). Traditionally, assessment of the patellar trajectory is subjectively performed by the surgeon during the operation, relying on direct visualization (Best et al. 2020). After placing the implants in the corresponding bones, the surgeon manually flexes and extends the knee of the anesthetized patient, to observe the range of motion of the joint and evaluate the patellar trajectory after the applied treatment. During this routine maneuver, the surgeon looks for any signs of lateral subluxation (movement of the patella to the outside of the knee) or malalignment. While the motion may appear simple because the patient lies on his back, without muscle activity due to anesthesia, this method showed to be relevant for assessing patellar tracking during TKR.

In this study, the maneuver has been replicated and two manual knee flexions and extensions (Figure 3) were performed to observe the patellar trajectory. The position of the optical markers was recorded using 18 infrared cameras (OptiTrack FLEX 3, Natural Point, Corvallis, OR, USA) at a sampling frequency of 100 Hz. Additionally, spring tensions were recorded using two tension load cells (RB-Phi-119, Phidgets, Calgary, Canada), and the pressure of the prosthetic button on the femur was measured using a compact pressure load cell (FX29, TE Connectivity, Wört, Germany), also at a sampling frequency of 100 Hz. A second-order Butterworth filter with a cutoff frequency of 12 Hz was applied to the optically captured marker trajectories (Javier Cuadrado et al. 2021), and a singular spectrum analysis (SSA) (Romero et al. 2015) with a window length of 30 was applied to the force measurements.

**Figure 3:**
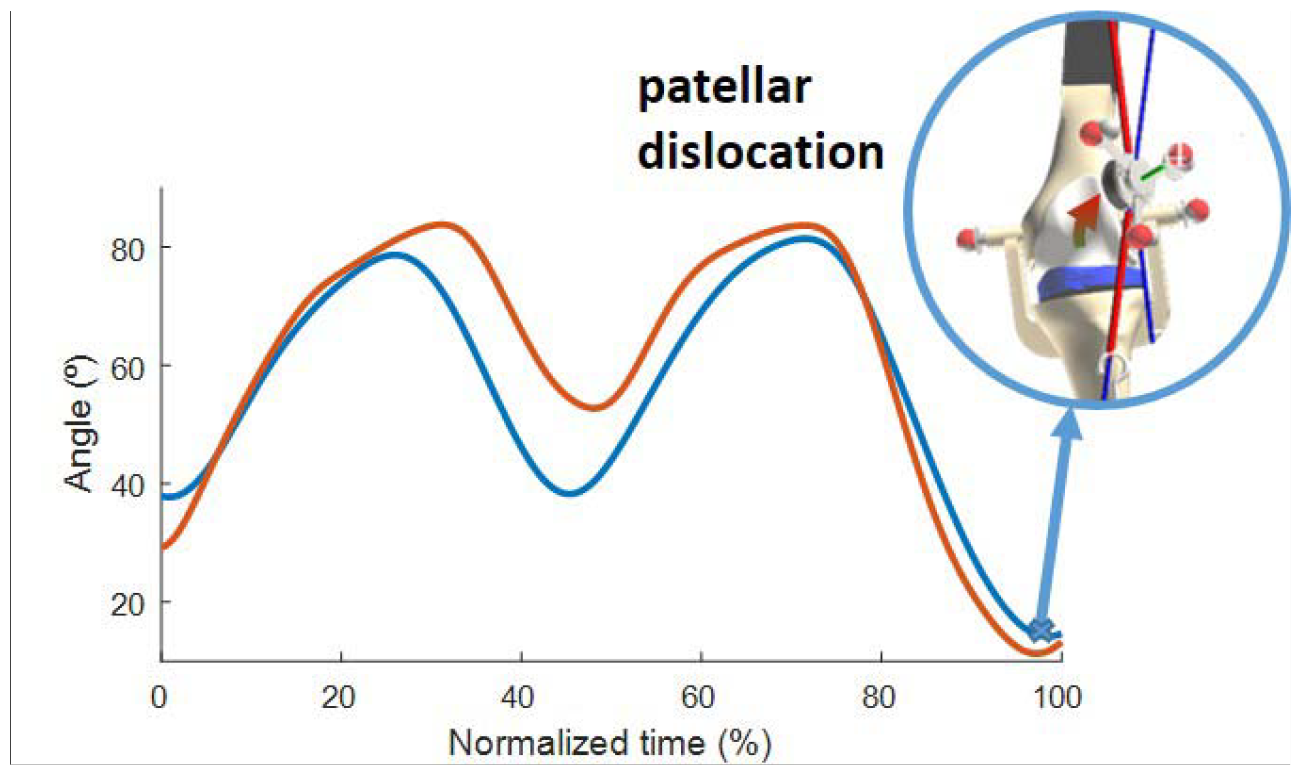
Knee angle during motion (blue: configuration A; orange: configuration B).

To validate different configurations, two distinct trajectories of the patella were measured by modifying the attachment point of the spring to the femur using the adjustable support (Figure 1). This modification corresponds to altering the Q angle, also known as the quadriceps angle, which measures the alignment of the quadriceps muscles and the patella relative to the femur (Khasawneh, Allouh, and Abu-El-Rub 2019). In this case, configuration A was laterally displaced by 20 mm compared to configuration B, resulting in a 4.55º difference between the respective Q angles. In configuration A, poor patellar tracking with patellar dislocation was generated when the knee flexion was lower than 20º (Figure 3). As shown in Figure 3, the motion started with the leg flexed around 35º, and was then flexed until 90º, extended to 45º, flexed again to 90º, and, finally, extended until 10º, thus resulting in a patellar dislocation for configuration A, but not for configuration B. Due to the manually executed experimental actuation, the imposed motion was not exactly the same for both configurations.

### 2.3. Computational Model

The leg model considered in this work consisted of three rigid bodies: the femur, the tibia-foot assembly, and the patella. The 3D geometries were identical to the physical pieces, both for the supports and the bones and implants. While the femur was fixed at the hip joint and could rotate about three directions, the joint between the femur and tibia was modeled as a hinge around the mechanical axis identified by two optical markers located on the sides of the knee (Figure 4, in purple). Thus, both the real leg mechanism and the virtual one had four degrees of freedom: three at the hip and one at the knee. The patella, on the other hand, was a free body with its six degrees of freedom, interacting with the leg through two linear springs-dampers (Figure 4, red lines: one attached to the femur and the other to the tibia) and the contact with the femoral implant.

**Figure 4:**
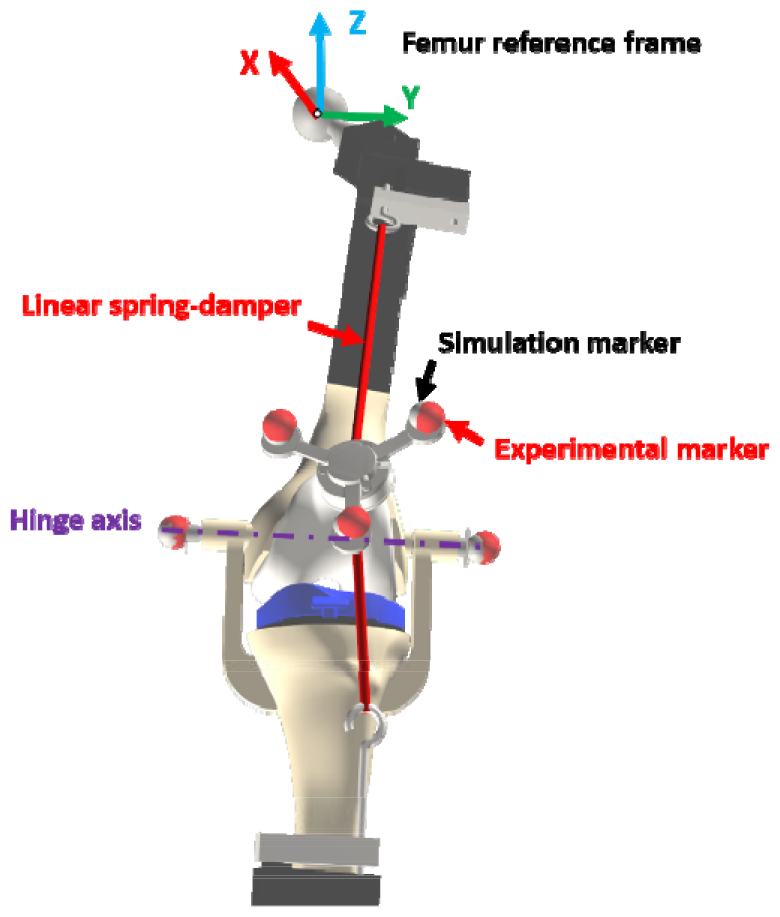
Computational model.

The geometrical and physical parameters of the rigid bodies (local coordinates of points, masses, inertias, etc.) were estimated from CAD models created in Solidworks and introduced, along with the mechanical constraints of the system, into a custom-developed library (Dopico 2016). The mechanical parameters of the springs (stiffness, natural length, and damping) were estimated from the experimentally recorded positions and forces during calibration.

### 2.4. Simulation

The primary objective of this work is to provide an affordable solution for experimental validation that can be replicated by research groups without access to extensive financial, experimental, and clinical resources. While this work presents a single simulation example, the authors’ future studies will compare different models and algorithms. This comparison will enable them to select the most accurate one based on experimental measurements of real-world phenomena.

#### 2.4.1 Guiding

The positions and orientations of the rigid bodies were obtained from the recorded marker positions captured by the cameras. To achieve this, the traditional approach described by Vaughan (Vaughan, Davis, and O’Connor 1999) was used, which consists of: (i) selecting three non-collinear entities, which can be markers or already located joints, within each segment; (ii) defining an orthogonal reference frame for the corresponding segment based on the three selected entities; (iii) using correlation equations to estimate the position and orientation of the rigid body (Javier Cuadrado et al. 2021).

The movements of the femur and tibia recorded by the motion capture optical system were used as inputs for the simulation. Therefore, the four free angles of the leg (three at the hip and one at the knee) were guided with the experimental values. The recorded movements of the patella served to experimentally validate the simulation results (Figure 4, red markers), as well as to approximate the patella to its initial static equilibrium position, which had to be determined.

#### 2.4.2 Formulation

The selected formulation for the multibody system dynamics in this work was the ALI3-P formulation, explained in (Dopico et al. 2014), which has been developed over many years as an evolution of the formulations presented in (J. Cuadrado et al. 2001; Bayo and Ledesma 1996). It is an Augmented Lagrangian formulation of index 3 in mixed coordinates (natural plus relative), with velocity and acceleration projections on the constraint subspaces.

The configuration of the multibody system was described by a set of *n*_*c*_ dependent coordinates 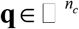. These coordinates were related through a set of *m* holonomic constraint equations **Φ**(**q**,*t*) = 0, and the equations of motion were expressed in the following manner:

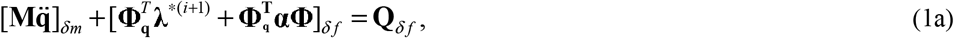

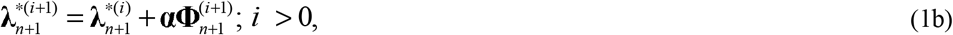

with **M = M(q)** and 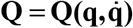 being the mass matrix and generalized force vector, respectively, **Φ**_**q**_ the Jacobian matrix of the constraints vector, λ the Lagrange multipliers vector, α a diagonal matrix that contains the penalty factors associated with the constraints, *δf* and *δm* scalar parameters of the generalized-α method, *n* the time step index, and *i* the iteration index of the approximate Lagrange multipliers’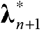. For a complete description of the formulation of the equations of motion and projections of velocities and accelerations, the reader is referred to (Dopico et al. 2019).

The integration scheme adopted was the Newmark integrator (Gavrea, Negrut, and Potra 2005), with a time-step size of 1 ms.

#### 2.4.3 Static Equilibrium

To perform a dynamics simulation of a multibody system, it is necessary to obtain a set of initial positions and velocities that satisfy the constraint equations, both at position and velocity levels. Additionally, in those multibody systems that have a defined static equilibrium position, it is advisable to start the simulation from the static equilibrium position to avoid the presence of initial high accelerations which could disturb the stability of the simulation. This involves solving the static equilibrium equations of the system to determine the equilibrium position. Unfortunately, when the system involves bodies in contact, solving the static equilibrium problem becomes very complex and there may even be multiple solutions.

In this work, the static equilibrium equations were obtained by eliminating the accelerations and velocity-dependent forces in the equations of motion, that is, by solving

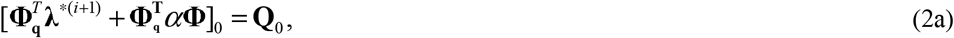

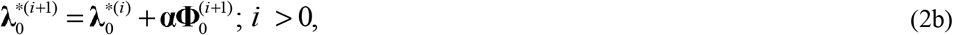

where i = 0, 1, 2,…, and the subscript 0 in the expressions of (1) indicates that quantities are evaluated at the initial time.

For solving the nonlinear system (1), a Newton-Raphson iteration was used, similar to the one used to solve the equations of motion (Dopico et al. 2019).

#### 2.4.4 Contact Model and Detection

Since the contact area had been lubricated in the test bench, the approach proposed in this work to address the contact problem between the patella and the femoral implant considers only the normal forces, and not the tangential forces (friction). The chosen normal force model for this work was the Flores model (Flores et al. 2011) and the expression for the normal force has the following form,

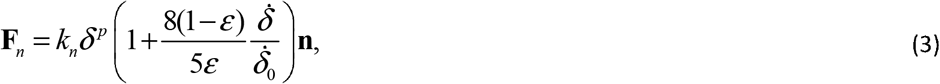

where *k*_*n*_ is the equivalent stiffness of the contact, which depends on the shape and the material properties of the colliding bodies, *p* is the Hertz’s exponent, *δ* is the indentation and 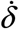 its temporal derivative,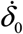 is the relative normal velocity between the colliding bodies when the contact is detected, *ε* is the coefficient of restitution, and **n** is the direction of the force. The subscript “**n**” comes from “normal”.

Since the colliding bodies had complex 3D geometries, a general collision detection algorithm was required. While a (triangular) mesh-to-mesh contact algorithm was used in previous works (Dopico et al. 2011, 2019), to check the penetration between bodies and find the corresponding contact points, in this work an algorithm based on the analytic formulation of the 3D geometry of the patellar prosthetic button was preferred (Yastrebov and Breitkopf 2013).

Analytic formulation involves using mathematical equations to calculate the distances between the surfaces of the geometric primitives. The specific equations used depend on the types of primitives involved. In this work, only the contact between the patellar button and the femur implant was considered. A sphere primitive was employed for the patellar button, since only its spherical portion would come into contact with the femur. A similar simplification was applied to the femur implant geometry, focusing solely on the surface that would interact with the patellar component (highlighted in orange in Figure 5a). Subsequently, this complex surface was approximated using a custom-made Matlab program by means of a 4^th^ order polynomial equation from the vertex coordinates with an R-squared value of 99.2% (Figure 5b).

**Figure 5:**
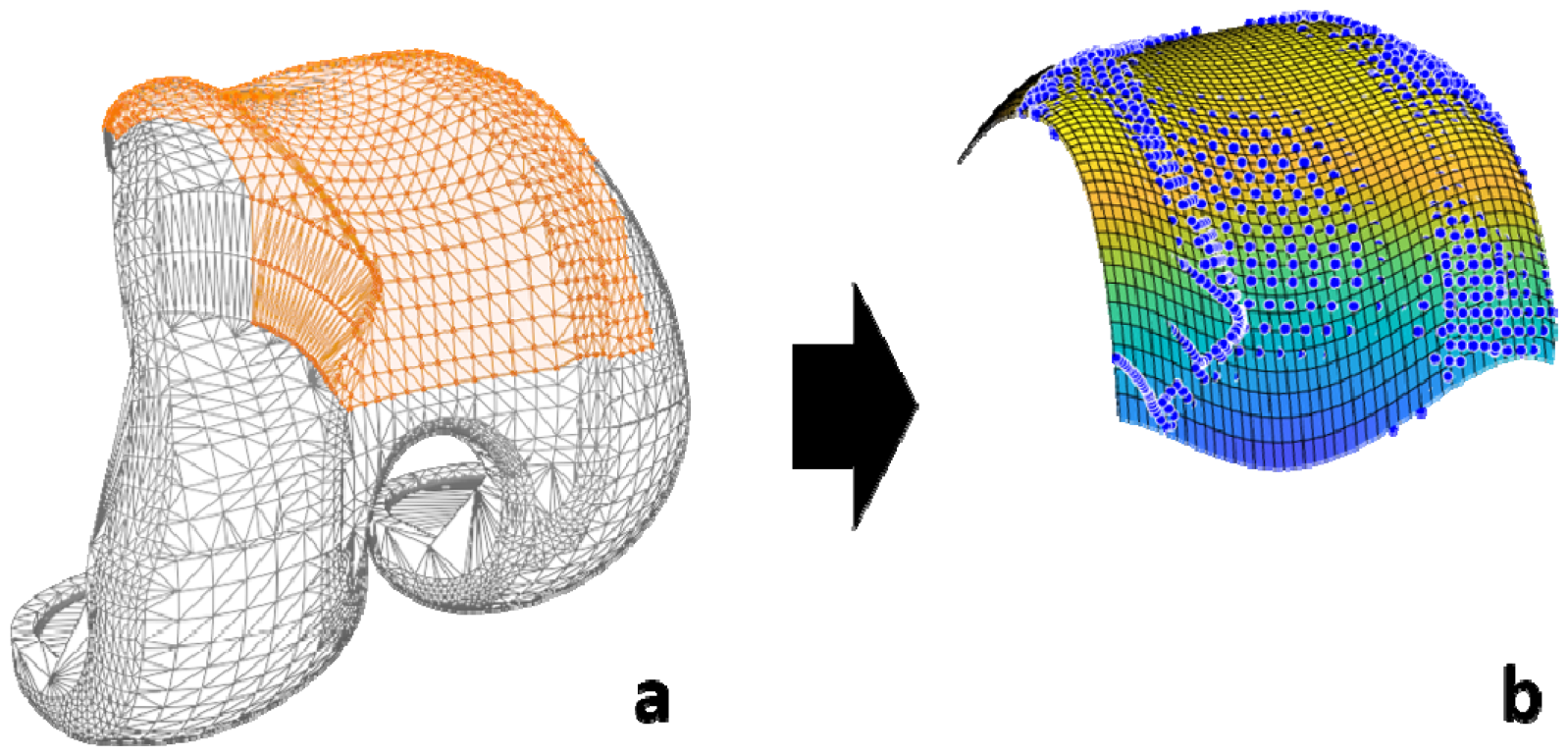
a) Contact surface considered; b) Analytical representation of the 3D geometry of the femur implant.

These equations take into account the position, orientation and size of the primitives. The simulation needs to determine whether or not two objects are in contact at a certain time-point. If the distance between the center of the sphere (patelar button) and the femur surface is smaller than the sphere radius, a collision or contact exists. Based on this information, and for each detected contact, the normal force is computed (perpendicular to the contact surfaces, Figure 6, in red) using the aforementioned contact model. The collision detection algorithm and the contact force model were implemented in the in-house developed library (Dopico 2016).

**Figure 6:**
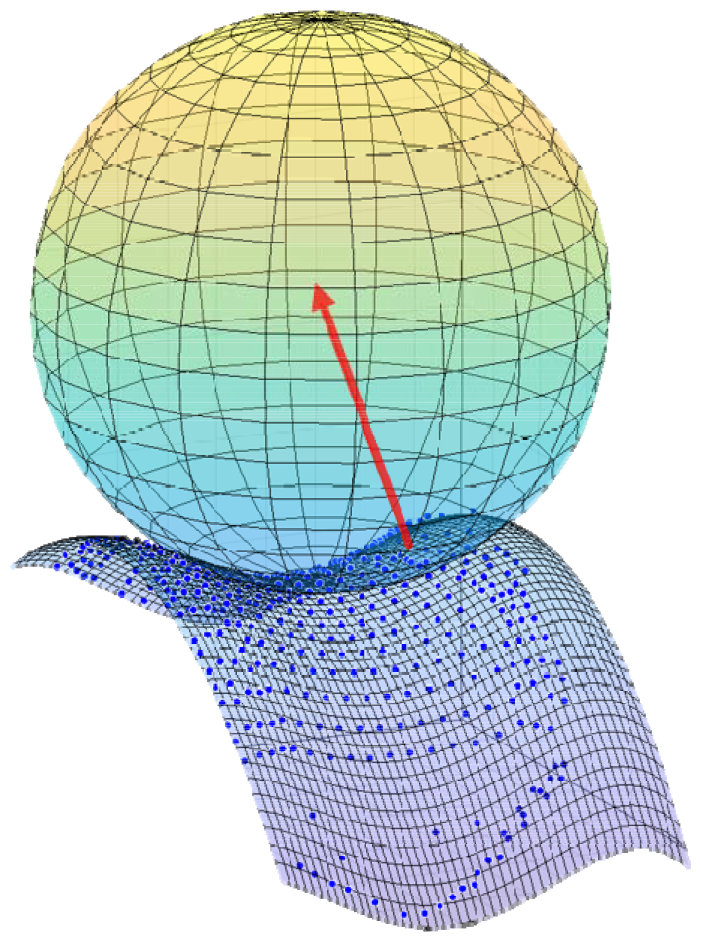
Representation of the normal contact force (red arrow) between the patellar prosthetic button and the femoral implant.

#### 2.4.5 Experimental Validation

In order to validate the results obtained from the computational simulations, the recorded experimental measurements were used as reference. The forces applied on the patella were validated by comparing the forces of the springs with the measurements from the load cells, and the contact force (only the normal component) with that obtained from the pressure sensor. The position of the patella during the motion was also validated by comparing the coordinates of the center of the patellar prosthetic button with the coordinates recorded by the optical motion capture system, both with respect to the femur reference frame to avoid error accumulation (Figure 4). The anatomical coordinate system of the femur was defined as follows: the origin was the hip center, the Y-axis was defined by the two markers located on each side of the knee, and the Z-axis was defined by the vector normal to the Y-axis and the vector between hip and knee centers. The error was quantified for the two configurations (A and B) by calculating the root mean square error (RMSE) between pairs of data sets.

#### 2.4.6 Computational details

All the analyses were run on an Intel(R) Core(TM) i7-13700KF @ 3.40 GHz, 32 GB RAM, SSD 2TB with operating system Windows 10 Pro. The single-threaded program was written in Fortran 2008 and C++, and compiled with MSVC 2017 and Intel Fortran 2018. The MA28 suit was used as the linear algebra package. To measure efficiency, the run-time was chosen, defined as the time required to solve the initial static equilibrium and run the simulation.

#### 2.4.7 Optimization

For both configurations, the results were first obtained using the spring parameters derived from calibration measurements. Subsequently, an optimization process was performed to enhance the results and observe their sensitivity to these parameters. The spring parameters were allowed to vary by 10% from their default values, and the objective function was defined as the sum of the RMSE of the forces (contact and springs) and the distance error of the relative position of the patella. The minimum value of the function was estimated in Matlab using the genetic algorithm (*ga* function).

## 3. Results

The authors successfully designed and built a low-cost, sensorized knee model using 3D printing for experimental validation of patellar tracking and contact simulation after TKR. This development provides an affordable solution for experimental validation that can be replicated by research groups with limited financial, experimental, and clinical resources. The total cost of the sensorized system, including the commercial training station, did not exceed 5000€ (motion capture system and 3D printer excluded). The recorded motion and force sensors enable the reproduction of movement in the virtual model, adjustment of the numerous simulation parameters, and experimental validation of the results. In addition, this work introduces a novel design of a sensorized patella by incorporating a pressure sensor.

In the case of the described simulation as validation example, the run-times required to simulate the 3.64 s of real motion of configuration A, and the 5.00 s of real motion of configuration B, were, respectively, 1.42 s and 1.67 s. This means, approximately, 2.77 times faster than real time.

The results indicate that the forces obtained through computer simulation (Figure 7; blue: original; yellow: optimized) followed a similar pattern to the forces obtained from experimental measurements (Figure 7, red). Despite the calibration, the forces from the springs at the initial instant (slightly bent knee) were approximated with a small error (less than 1.5 N error for the configuration A and 4.2 N for configuration B, without optimization). During knee flexion, the forces of the springs increased and generated more pressure on the patella. However, the tibial spring force did not increase throughout the complete flexion. From a certain angle of knee flexion, both configurations showed a reduction in spring length and force. This behavior was accurately replicated by the simulation.

**Figure 7:**
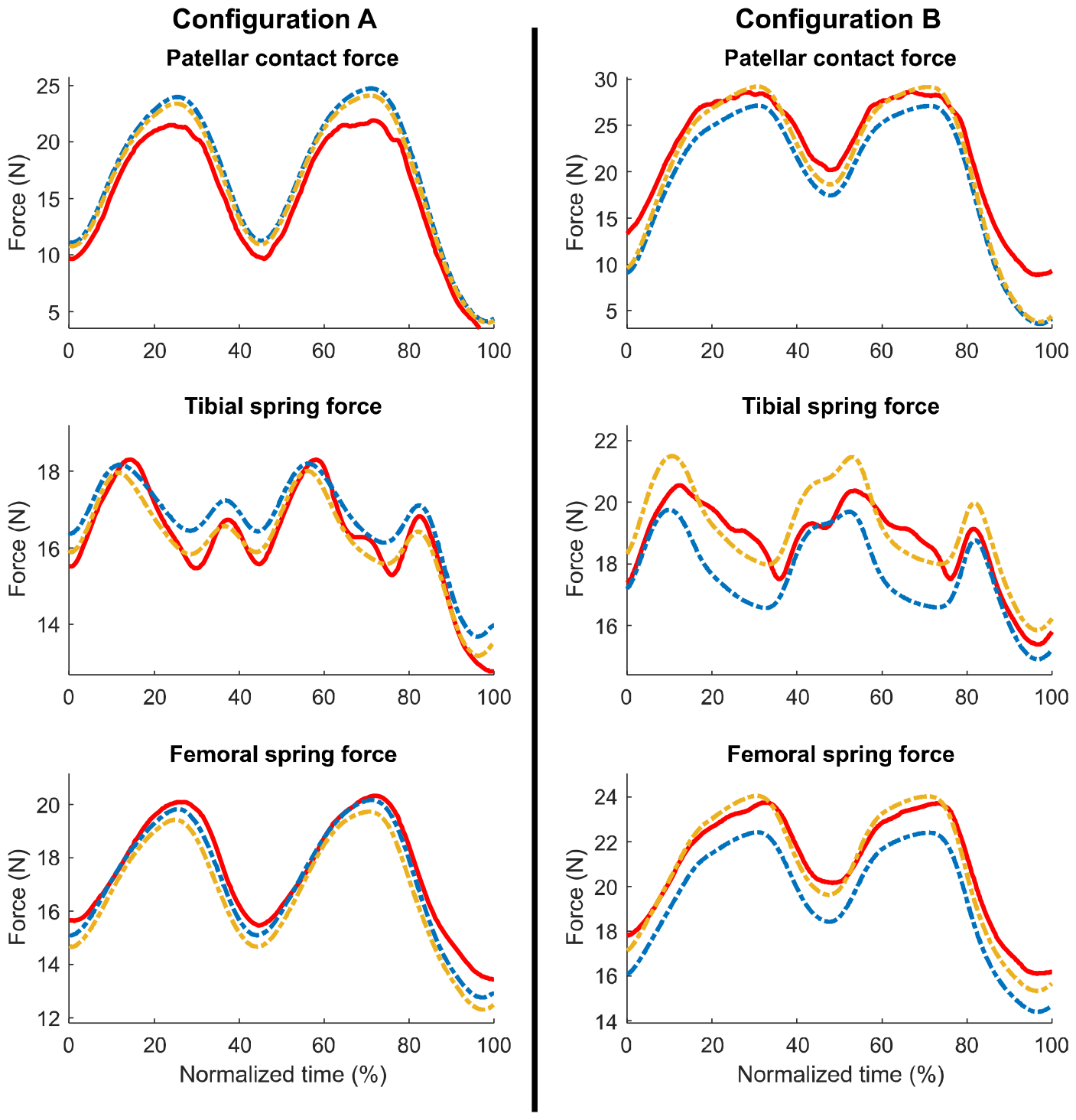
Comparison between experimental forces (red) and simulation forces (blue: original; yellow: optimized).

The peak values of the forces show a proportional error to the initial offset, since it was maintained throughout the motion. The highest error was observed for the contact force in configuration A, with errors of 2.8 N and 2.2 N before and after optimization, respectively. Regarding the overall simulation, the forces were estimated with high accuracy. The mean of the RMSEs of all forces were below 2 N with the default parameters, and below 1.2 N after optimization (Table 1).

**Table 1:**
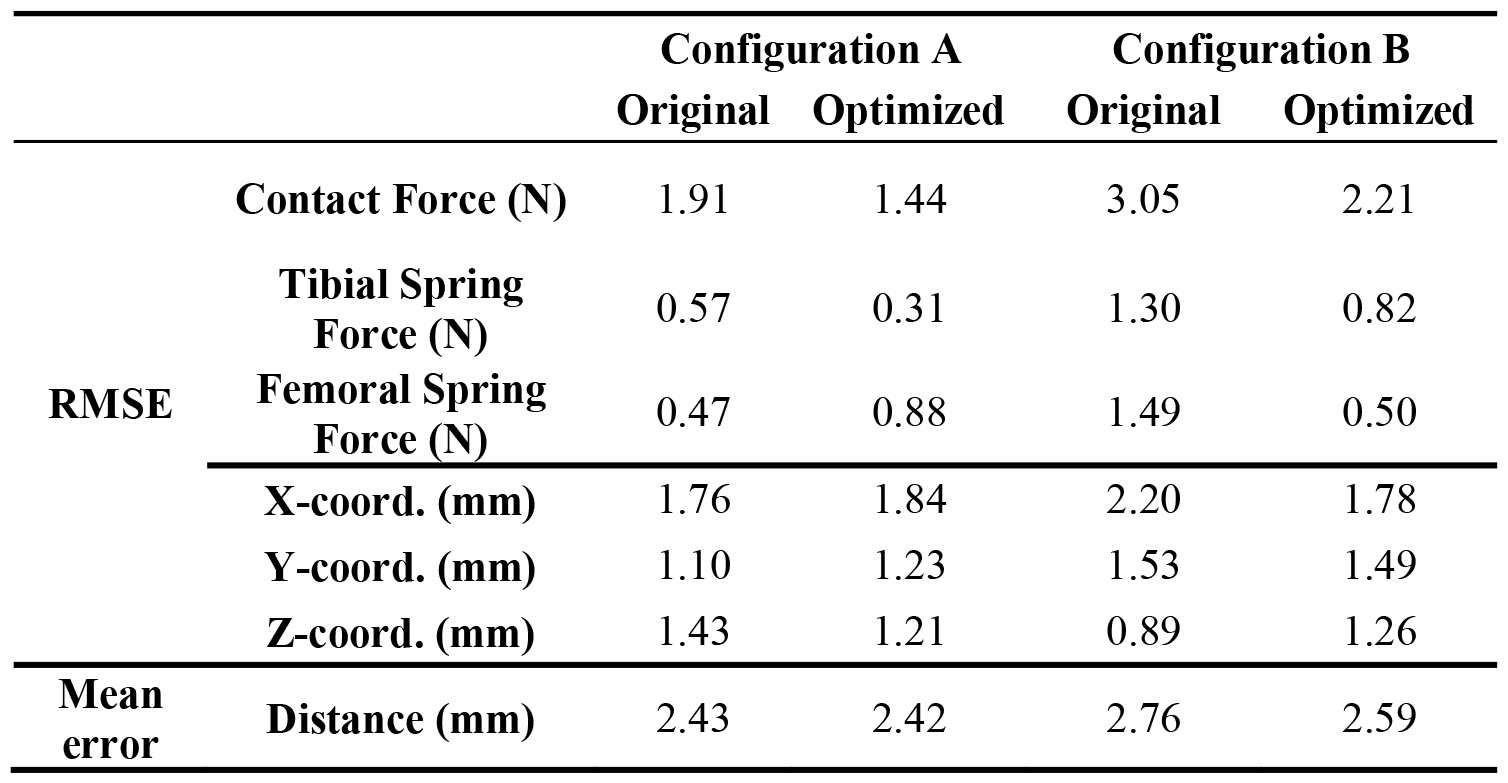
Comparison of simulation errors with respect to experimental data.

The simulated movement of the patella also showed very small discrepancies with respect to the motion recorded by the optical motion capture system (Figure 8). It was observed that the patella started its movement with positional errors of 3.1 mm and 3.2 mm for configurations A and B, respectively, resulting in discrepancies in all the three coordinates of the studied point. The average error incurred during the complete motion was lower than 2.8 mm for distance, and lower than 2.2 mm for the coordinates, consistent across all the simulations (Table 1). Patellar dislocation was successfully reproduced during the simulation of configuration A, both with the optimized and the default values of the parameters. While all force estimations showed some improvement in the simulations with optimized parameters, the patellar motion remained almost unchanged.

**Figure 8:**
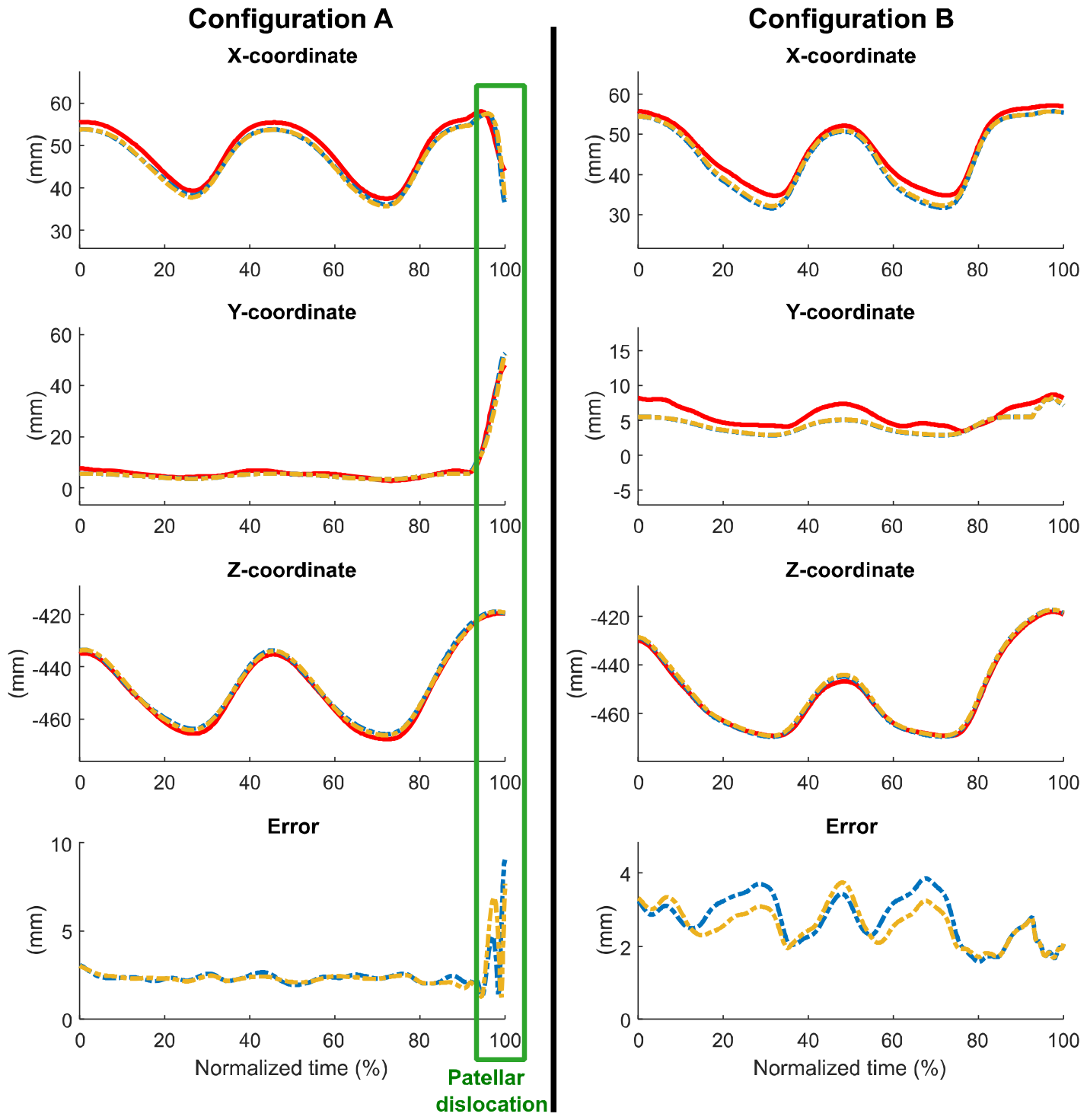
Comparison of experimental (red) and computational patellar positions (blue: original, yellow: optimized).

## 4. Discussion

In this study, a commercial surgical training station was transformed into a sensorized test bench to provide an affordable solution for experimental validation that can be replicated by research groups without access to extensive financial, experimental, and clinical resources. The use of 3D-printed models and sensors allowed low-cost and reproducible experimental validation of computer simulation (patellar movement and forces) while avoiding ethical issues. The cost of the proposed system is very affordable compared to knee simulators developed by universities like Oxford, Kansas, and Purdue (Maletsky and Hillberry 2005; Halloran et al. 2010). Half of the total prize was due to the purchase of the commercial training station, which could be replaced by simpler mechanical supports. Furthermore, in this work, a novel fully sensorized patella has been designed by incorporating a pressure sensor, which has the potential to lead to the development of new instrumented implants or to improvements in existing knee simulators.

Before attempting to predict treatments, the objective of this preliminary study was to demonstrate the ability to faithfully replicate the surgeon’s maneuver, which has clinical relevance. While this common maneuver may seem simple, it is sensitive enough to detect signs of lateral subluxation (movement of the patella to the outside of the knee) or malalignment. It has been demonstrated that, by making minor adjustments to the Q angle, a patellar dislocation can be observed.

The authors acknowledge that the knee joint is more complex than a simple revolute joint. However, this complexity is not relevant to the patellar trajectory studied in this work, which is defined as the movement of the kneecap relative to the femoral groove during knee flexion and extension. Since the motion of the tibia and femur was controlled in this work, the joint linking them becomes irrelevant. Even if the tibia were defined as an independent body, it would only affect to the position of the attached point of the spring in the experimental motion capture and the simulation. This study demonstrated that changes resulting from the modification of the attachment points of the springs were accurately reproduced. Additionally, it must be noted that, the surgeon’s maneuver, as well as the replicated experimental measurement, involve manual actuation. This makes it very difficult to apply the corresponding forces to the virtual model since the force magnitude and the application points/surfaces are not evident. This issue needs to be resolved, especially if the contact between the femur and tibia implants is added to the problem for a more comprehensive treatment prediction.

Although the system was simplified for this preliminary study, the mechanical behavior exhibited was quite complex due to the contacts between bodies, resulting in a complex solution for the initial static equilibrium position, including the existence of multiple equilibrium positions (Dopico et al. 2019). The obtained results showed that the computationally simulated forces followed patterns similar to those of the experimentally measured forces, but the initial offsets were maintained throughout the motion. These differences could be the result of small discrepancies in the equilibrium position and slight discrepancies in the spring parameters. However, force predictions were good and were improved by optimizing the spring parameters. The overestimation of patellar contact forces in configuration A could be attributed to an underestimation of the measurements, caused by the actuation of non-centered force pressures on the sensor. A slight difference was also observed in the initial position of the patella compared to the position recorded by the optical motion capture system. When attempting to position the patella at the experimentally recorded initial position, it was observed that the 3D geometries of the patella and femoral implant were not in contact, but rather had a small separation of some few millimeters. This could be due to defects in 3D printing, inaccuracies in optical measurements exacerbated by their processing and the body motion reconstruction from them, or inaccuracies in the analytical approximation of the geometry of the femur implant. While the optical motion capture system is considered a reference in terms of precision (Liao et al. 2020; Javier Cuadrado et al. 2021), it is quite reasonable to expect errors of a couple of millimeters within a capture volume of more than 25 m^3^ with markers of 14 mm in diameter.

As the knee flexed, the spring forces increased, exerting greater pressure on the patella. During extension, the forces decreased, and, in configuration A, the observed patellar dislocation along the second extension was successfully reproduced, even when using the default values of spring parameters.

Despite the small discrepancies, which are common between the real and virtual worlds, the obtained results were very satisfactory and allowed validation of the models used in the study.

These results will also pave the way for other implementations (e.g., for other treatments) and will serve to communicate with healthcare professionals.

As future work, more tests will be conducted to simulate new configurations, by modifying the orientation of the hinge axis and the insertions of tendons. Furthermore, the detail of the system will be increased in order to better reflect anatomy, including the use of springs with properties that are more similar to those of tendons, and consideration of the contact between tibia and femur, among other aspects. In order to assess the impact of changes in the system parameters, and to improve the stability and robustness of the simulations, a sensitivity analysis will be conducted. Additionally, a (triangular) mesh-to-mesh contact algorithm will be implemented to compare its efficiency and accuracy with the currently implemented analytical algorithm.

Finally, it is worth mentioning that, while the combination of 3D-printed models and sensors can be used to obtain measurements of physically realistic representations of bone pathologies and treatments, computer simulation proves to be a much more cost-effective and faster tool to use, once implemented.

## 5. Conclusions

The authors have presented a low-cost and reproducible knee simulator that avoids ethical issues by utilizing 3D-printed models and sensors. This model serves as a valuable tool for validating patellar tracking and contact simulation post-TKR, offering a cost-effective solution accessible to research groups with restricted financial, experimental, and clinical resources. With the aid of recorded motion and force sensors, the system allows the recreation of virtual movements, fine-tuning of numerous simulation parameters, and validation of experimental outcomes. Furthermore, this study introduces an innovative sensorized patella design, featuring a pressure sensor integration.

## Availability of supporting data

The datasets generated for this study are available on request to the corresponding author.

## Ethical Approval and Consent to participate

Not applicable.

## Authors’ contribution

F.Mi. designed the experiments with the supervision of J.C. F.Mi. performed the experiments. F.Mo. implemented the contact algorithm in the in-house library developed by D.D. F.Mi. analyzed the data. F.Mi. and J.C. wrote the manuscript in consultation with D.D. and F.Mo.

